# Fine spectral tuning of a flavin-binding fluorescent protein for multicolor imaging

**DOI:** 10.1101/2022.12.09.519645

**Authors:** Andrey Nikolaev, Anna Yudenko, Anastasia Smolentseva, Andrey Bogorodskiy, Fedor Tsybrov, Valentin Borshchevskiy, Siarhei Bukhalovich, Vera V. Nazarenko, Elizaveta Kuznetsova, Oleg Semenov, Alina Remeeva, Ivan Gushchin

## Abstract

Flavin-binding fluorescent proteins (FbFPs) are promising genetically encoded tags for microscopy. However, spectral properties of their chromophores (riboflavin, flavin mononucleotide and flavin adenine dinucleotide) are notoriously similar even between different protein families, which limits applications of flavoproteins in multicolor imaging. Here, we present a palette of twenty-two finely tuned fluorescent tags based on the thermostable LOV domain from *Chloroflexus aggregans* (CagFbFP). We performed site saturation mutagenesis of three amino acid positions in the flavin-binding pocket, including the photoactive cysteine, to obtain variants with fluorescence emission maxima uniformly covering the wavelength range from 486 to 512 nm. We demonstrate three-color imaging based on spectral separation and two-color fluorescence lifetime imaging using the proteins from the palette. These results highlight the possibility of fine spectral tuning of flavoproteins and pave the way for further applications of FbFPs in fluorescence microscopy.

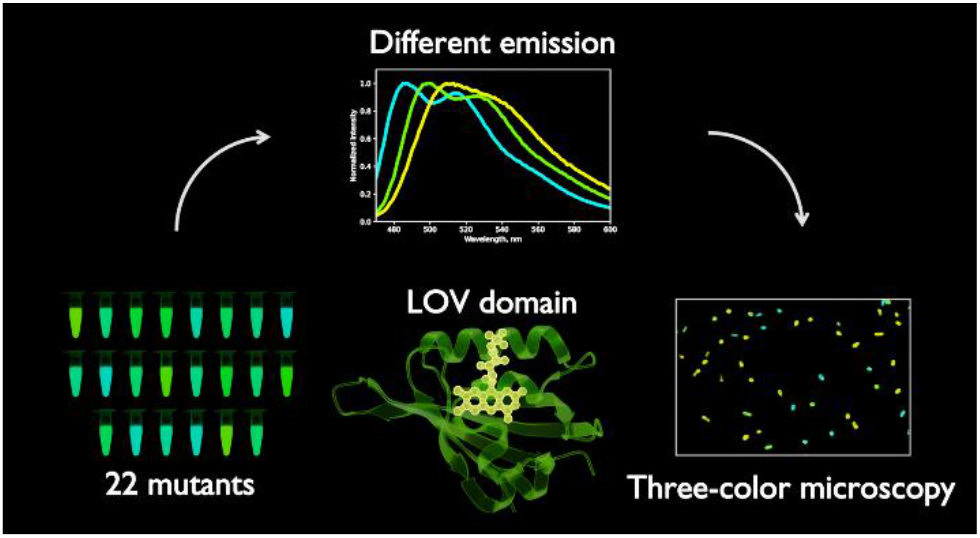

## Introduction

Flavins, such as riboflavin (RF, vitamin B_2_), flavin mononucleotide (FMN) and flavin adenine dinucleotide (FAD), are ubiquitous biomolecules present in virtually all living cells (1, 2). They are often employed as cofactors by so-called flavoproteins, whose genes constitute a notable fraction of all genes found in the genomes of variable organisms (3, 4). On their own, flavins display complex photochemistry and photophysics, and have been the focus of numerous studies (5–8). However, while the spectra of other biological chromophores, such as retinal (9–11) or tetrapyrroles (12–14), are strongly influenced by their protein environment, the positions of absorption and fluorescence maxima of RF, FMN and FAD are vastly similar between different protein families (5, 6). Several studies have suggested the routes for spectral tuning of flavins (15–17), yet at the moment flavin-binding proteins have not been designed to exhibit pronounced bathochromic or hypsochromic spectral shifts.

Among the different families of flavoproteins, LOV domains found especially many applications as molecular biology tools (12, 18). LOVs are sensor modules of natural photosensitive proteins found in plants, fungi, bacteria, archaea, and protists (19). They absorb UV and blue light via noncovalently bound FMN or FAD in oxidized state with the characteristic absorption peak around 450 nm. They may also bind RF, which results in altered thermal stability and photobleaching kinetics, but does not affect spectral properties (20, 21). Most of the natural LOVs have a conserved cysteine near the flavin isoalloxazine moiety. Following absorption of a photon, a covalent bond is formed between the cysteine and the flavin, and the protein becomes non-fluorescent until the ground state is restored. In engineered flavin-binding fluorescent proteins (FbFPs), the cysteine is replaced with a non-reactive amino acid such as alanine. Developed in 2007, FbFPs exhibit a number of desirable properties: (i) small gene (300-360 base pairs) and protein (10–12 kDa) size; (ii) no requirement for molecular oxygen or supplementation of exogenous chromophores; (iii) no need for chromophore maturation (22).

Generally, LOV domains are highly amenable to engineering. Several different strategies have been employed to obtain improved LOV-based molecular biology tools (23). For example, FbFPs with enhanced optical (24, 25) or thermodynamic (26, 27) properties were generated, as well as LOVs with altered switch-off kinetics (28–30), signal transduction mechanism (31) or dissociation constants (32).

At the same time, flavin-binding fluorescent proteins are poorly amenable to tuning of absorption and fluorescence spectra. Most of the mutations described to date lead to minor hypsochromic shifts, with the exception of those in the singlet oxygen generator miniSOG2 protein (33). Replacement of the conserved glutamine adjacent to FMN with a valine or a leucine results in a blue shift of ~10 nm (34–37), although a similar mutation in the phiSOG has almost no effect on the spectrum (36). Replacement of the glutamine with positively charged amino acids, contrary to the prediction (38), also resulted in hypsochromic shifts (39, 40). Stabilization of lysine side chain near the flavin leads to a red shift of 5-7 nm (41). Finally, mutation of flavin-adjacent asparagine can also lead to a 4-6 nm red shift (42, 43).

In this work, we set out to expand the range of color-tuned FbFPs and examine the possibility of fine tuning of flavin spectral properties. We tested the effects of random mutations of the conserved photoactive cysteine and two other amino acids on the properties of FbFP based on a thermostable LOV domain from *Chloroflexus aggregans* (CagFbFP). We obtained 22 variants with the emission maxima uniformly covering the spectral range from 486 nm (LOV domains with a glutamine to valine mutation) to 512 nm (observed for a double mutant I52V A85Q). To demonstrate the applicability of the resulting palette of FbFPs for multicolor imaging, we performed three-color imaging based on spectral separation and two-color fluorescence lifetime imaging with differentially labeled *E. coli* cultures.

## Results

### Shape and fitting of fluorescence spectra

FbFPs share a highly conservative fluorescence emission spectrum with two maxima at ~500 and ~525 nm, and a wide shoulder in the long wavelength region (Figure 1). Previously, while searching for color-tuned variants, we analyzed the differences between the positions of absolute fluorescence emission maxima of mutated and parent proteins. However, we also identified variants with significantly deformed spectra, where the two emission maxima merge into one (see Figure 2 for examples). For such variants, the nominal shift of the emission maximum does not correspond to the shift of the whole spectrum. Therefore, in addition to analyzing the position of the maximum, we also analyzed the perceived color of the emitted light (hue H in the HSV color representation), which reflects the overall position of the spectrum. For a more accurate determination of the position of maxima, we fitted the spectra with Gaussians (Figure 1).

**Figure 1.**
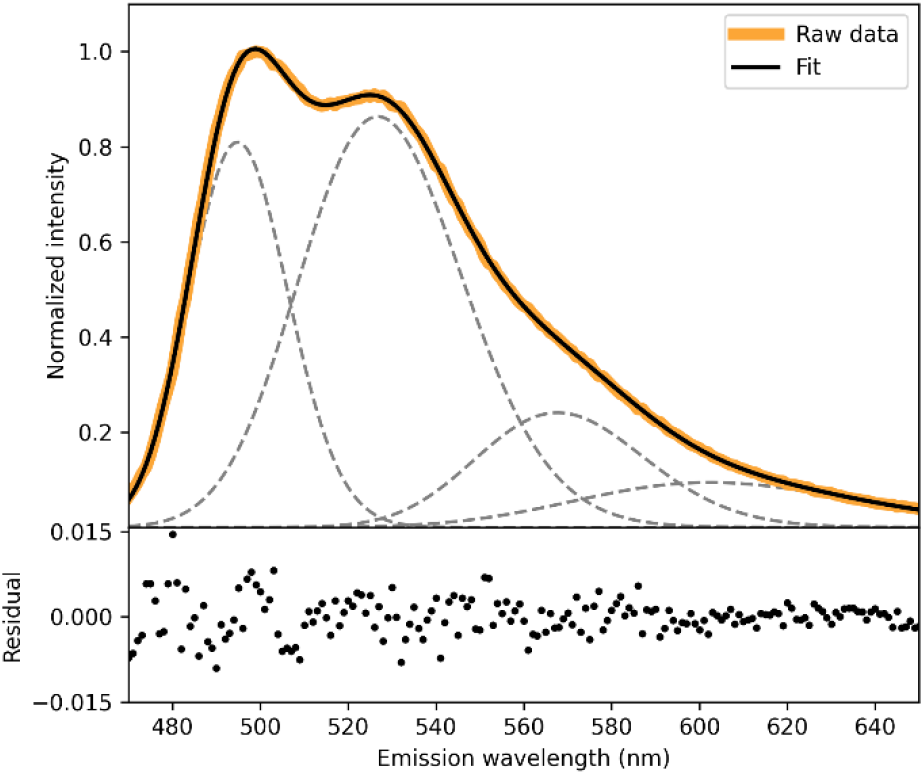
Fluorescence emission spectrum of CagFbFP and its fit as a sum of four Gaussians (gray dashed lines).

**Figure 2.**
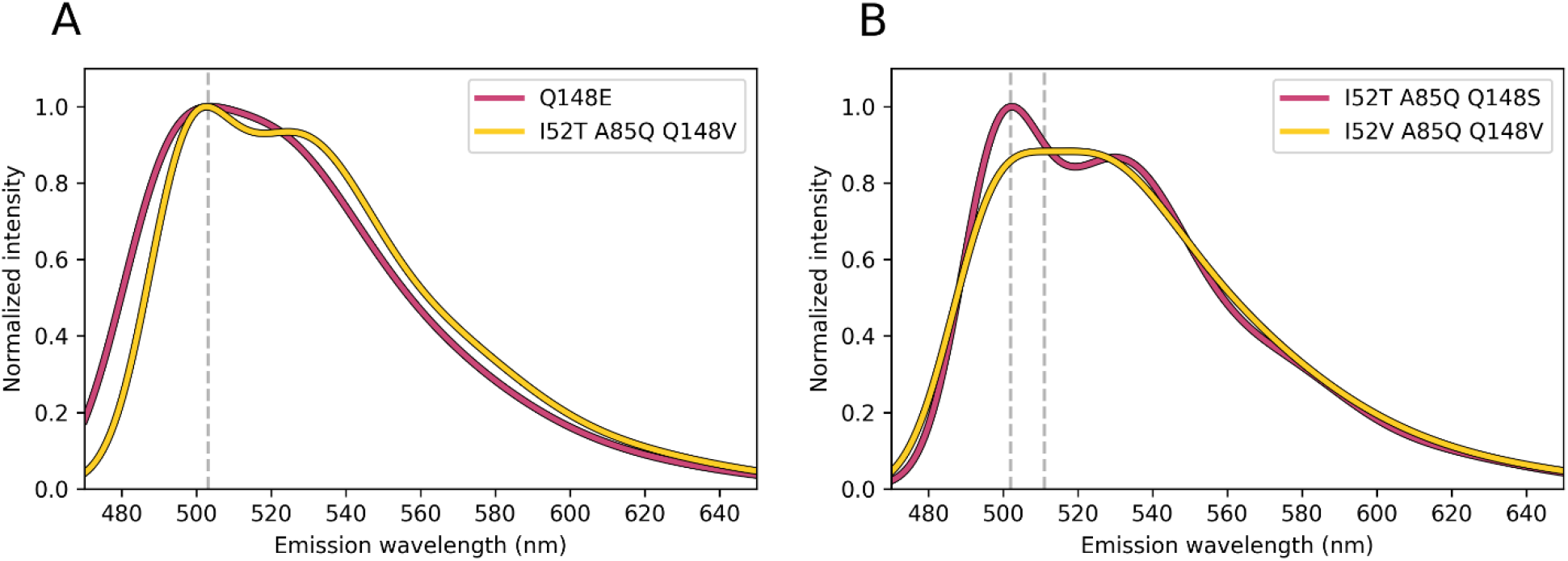
Examples of the fluorescence emission spectra of two CagFbFP variants with similar positions of the emission maxima but overall shifted spectra (A) and two variants with different nominal emission maxima but not shifted spectra (B). Positions of the emission maxima are denoted with dashed lines.

### Random mutagenesis of the photoactive cysteine position

Most of natural LOV domains contain a photoactive cysteine (19, 44). In engineered LOVs, this cysteine has been replaced with alanine, glycine, serine or proline. We reasoned that substituting it with some other amino acid may produce noticeable spectral alterations, given that the position is close to the flavin chromophore (Figure 3A). Intriguingly, the effects of other substitutions have not been reported previously, and we decided to conduct site saturation mutagenesis for this position. To have a higher chance of obtaining a folded protein even after very destabilizing mutations, we chose the recently developed thermostable fluorescent reporter CagFbFP (45) as a template. We found that the majority of the colonies with random mutations A85→X either did not display altered spectra or lost fluorescent properties. Only emission of the variant A85Q was red-shifted for 10 nm, yet the protein had extremely low expression level and stability (melting temperature of 42 °C, ~40 °C lower than that of CagFbFP).

**Figure 3.**
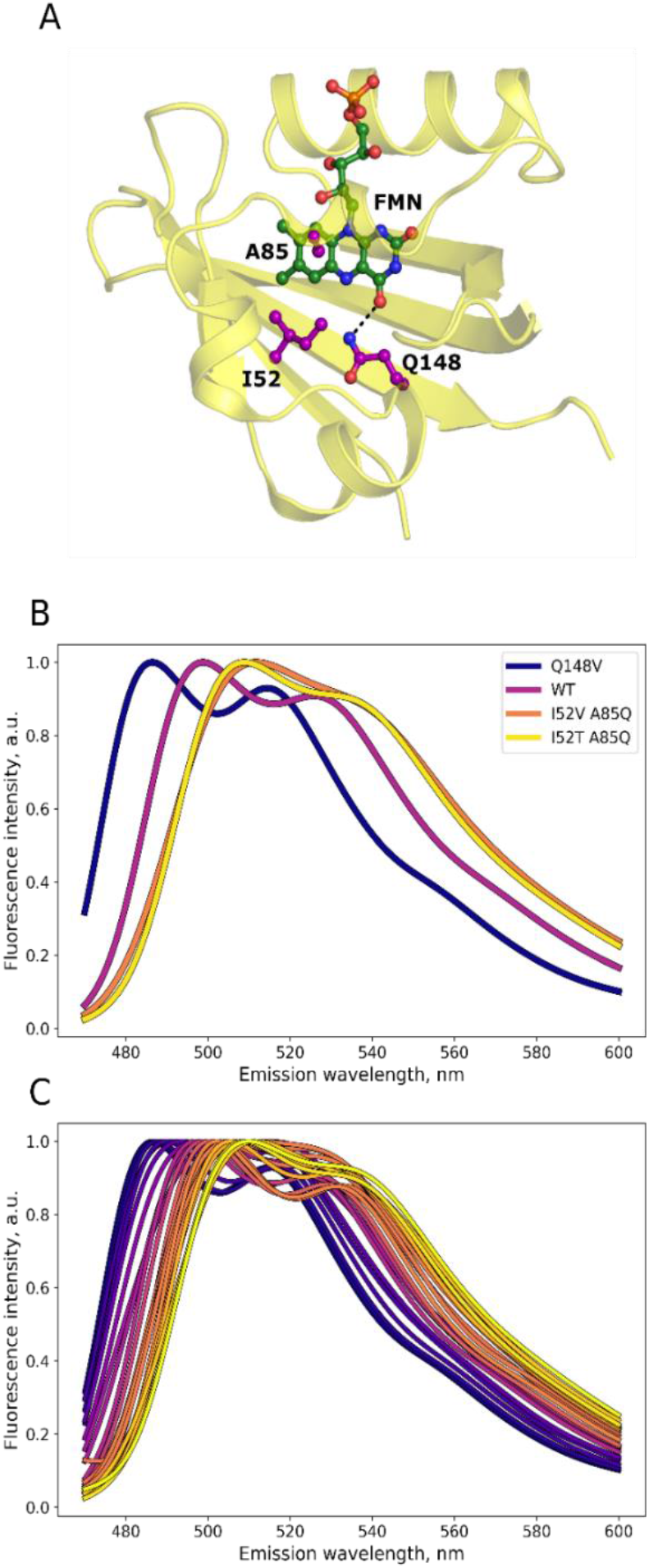
Palette of CagFbFP variants with altered fluorescence emission spectra. (A) Structural model of CagFbFP (45) with the side chains of mutated amino acids I52, A85 (corresponding to photoactive cysteine in natural LOV domains) and Q148 shown in purple. (B) Fluorescence emission spectra of the parent protein CagFbFP, the blue-shifted variant Q148V, and the red-shifted variants A85Q I52T and A85Q I52V. (C) Emission spectra of all identified CagFbFP variants (Table 1). The spectra are colored blue to yellow for ease of perception.

Given that the side chain of glutamine is bigger than that of native cysteine, we hypothesized that compensating mutations of the nearby I52 (Figure 3A) to a polar or smaller residue might stabilize the variant A85Q. First, we produced double mutants A85Q I52T and A85Q I52V. Both mutations significantly increased the expression level and raised the thermal stability by ~10 °C (Table 1). Site saturation mutagenesis I52→X performed on the A85Q variant produced mostly non-fluorescent cell colonies. Besides the mutations I52V and I52T, we identified the variant I52A A85Q, which has an untypical spectrum with the two emission peaks merging into one, along with low thermal stability (unfolding at 42 °C) and very low expression level.

**Table 1.** Main properties of the color-shifted CagFbFP variants.

| Hue | $\lambda_{\text{exc}}^1$<br>(nm) | $\lambda_{\text{em}}^1$<br>(nm) | $\lambda_{\text{fit1}}^1$<br>(nm) | $\lambda_{\text{fit2}}^1$<br>(nm) | Fluorescence<br>lifetime $\tau_{\text{av}}$, (ns)<sup>2</sup> | $T_{\text{m1}}$ (°C)<sup>2</sup> | $T_{\text{m2}}$ (°C)<sup>2</sup> | Expression<br>(mg/L) | Mutations |
| --- | --- | --- | --- | --- | --- | --- | --- | --- | --- |
| 154.1 | 443 | 486 | 483 | 513 | 3,76±0,05 | 71.4±0.1 | 81.1±0.3 | 108 | Q148V |
| 153.6 | 443 | 487 | 484 | 514 | 3,90±0,05 | 73.4±0.2 | 82.5±0.3 | 110 | Q148I |
| 152.7 | 444 | 488 | 484 | 515 | 3,93±0,04 | 71.7±0.1 | 81.0±0.2 | 112 | Q148L |
| 151.0 | 444 | 491 | 484 | 512 | 3,24±0,03 | 61.1±0.3 | 71.5±0.2<sup>3</sup> | 21 | Q148H |
| 151.0 | 445 | 490 | 485 | 516 | 3,98±0,13 | 69.3±0.5 | 79.0±0.3 | 19 | Q148M |
| 149.2 | 445 | 494 | 486 | 515 | 3,88±0,09 | 60.6±0.4 | 74.0±0.5 | 14 | Q148S |
| 146.1 | 448 | 498 | 488 | 515 | 4,19±0,07 | 47.2±0.8 | 59.0±0.1 | 20 | Q148R |
| 144.4 | 449 | 504 | 491 | 517 | 4,51±0,08 | 55.9±0.2 | 71.9±0.3 | 10 | Q148E |
| 143.8 | 448 | 499 | 491 | 523 | 4,33±0,10 | 59.7±0.4 | 73.2±0.5 | 12 | Q148G |
| 142.6 | 449 | 497 | 493 | 524 | 4,16±0,10 | 57.5±0.1 | 71.7±0.3 | 8 | I52V A85S |
| 140.4 | 450 | 499 | 494 | 526 | 4,53±0,03 | 64.3±0.2 | 78.1±0.5 | 50 | Original protein |
| 137.2 | 452 | 500 | 496 | 529 | 4,68±0,06 | 56.7±0.2 | 67.7±0.2 | 15 | I52V A85V |
| 135.8 | 450 | 507 | 495 | 524 | 3,74±0,12 |  | 55.6±0.2 | 8 | I52T A85Q Q148C |
| 135.7 | 452 | 503 | 496 | 527 | 4,26±0,06 | 46.3±0.2 | 58.9±0.1 | 11 | I52T A85Q Q148V |
| 135.0 | 454 | 503 | 499 | 526 | 3,66±0,07 |  | 55.8±0.2 | 5 | I52V A85Q Q148L |
| 134.6 | 450 | 511 | 496 | 521 | 3,67±0,17 |  | 55.0±0.2 | 12 | I52T A85Q Q148S |
| 134.6 | 451 | 506 | 497 | 526 | 3,86±0,10 |  | 52.4±0.9 | 9 | I52T A85Q Q148F |
| 134.0 | 455 | 502 | 498 | 531 | 4,40±0,05 | 49.6±1.0 | 58.4±0.8 | 34 | I52V A85Q Q148V |
| 131.5 | 453 | 508 | 500 | 531 | 4,24±0,06 | 45.9±0.5 | 59.0±0.3 | 12 | I52T A85Q Q148H |
| 130.5 | 454 | 506 | 500 | 533 | 4,30±0,11 |  | 47.4±0.4 | 9 | I52V A85Q Q148A |
| 129.3 | 452 | 512 | 503 | 536 | 4,57±0,07 |  | 48.6±1.3 | 13 | I52V A85Q |
| 127.1 | 455 | 509 | 502 | 534 | 4,65±0,06 | 44.9±0.1 | 56.7±0.1 | 7 | I52T A85Q |
| 125.8 | 454 | 510 | 502 | 533 | 4,54±0,07 |  | 42.1±0.1 | 8 | I52V A85Q Q148N |

<sup>1</sup> $\lambda_{\text{exc}}$ is the position of the excitation maximum, $\lambda_{\text{em}}$ is the position of the emission maximum, $\lambda_{\text{fit1}}$ and $\lambda_{\text{fit2}}$ are positions of the fitted Gaussians corresponding to the first and second peaks.
<sup>2</sup> The errors are the standard deviations in three independent experiments. $T_{\text{m1}}$ and $T_{\text{m2}}$ are the temperatures of the first and second melting transitions, correspondingly.
<sup>3</sup> The third transition is observed at 47.0±1.4 °C.

We also tested the effects of mutations A85→X on the I52V variant of CagFbFP. No new variants with significant shifts were observed; I52V A85Q was obtained again, I52V A85S was slightly blue-shifted and I52V A85V was slightly red-shifted compared to CagFbFP (Table 1).

### Random mutagenesis of the conserved glutamine

Previously, we tested the effects of substitutions of the conserved glutamine Q148 (Figure 3A) with polar and charged amino acids (40), all of which displayed hypsochromic shifts reaching ~6 nm. Here, we performed Q148→X random mutagenesis of CagFbFP and identified six new blue shifted variants with a maximum shift of 12 nm. Searching for more red-shifted variants, we also tested the effects of Q148→X mutagenesis on the already red-shifted variants I52V A85Q and I52T A85Q. Only one variant, I52T A85Q Q148N, gained an additional minor red shift, while 9 other mutants uniformly covered the range of spectral shifts between it and the original CagFbFP.

### Palette of finely tuned FbFPs

Based on the results of our site saturation mutagenesis rounds, we identified a palette of 22 variants, which, together with CagFbFP, uniformly cover the range of emission spectra with the maxima from 486 to 512 nm (Figure 3B, C and Table 1). The excitation spectra show less variation (Figure 4). We have also measured fluorescence lifetime and thermal stability of the identified variants *in vitro* (Table 1).

**Figure 4.**
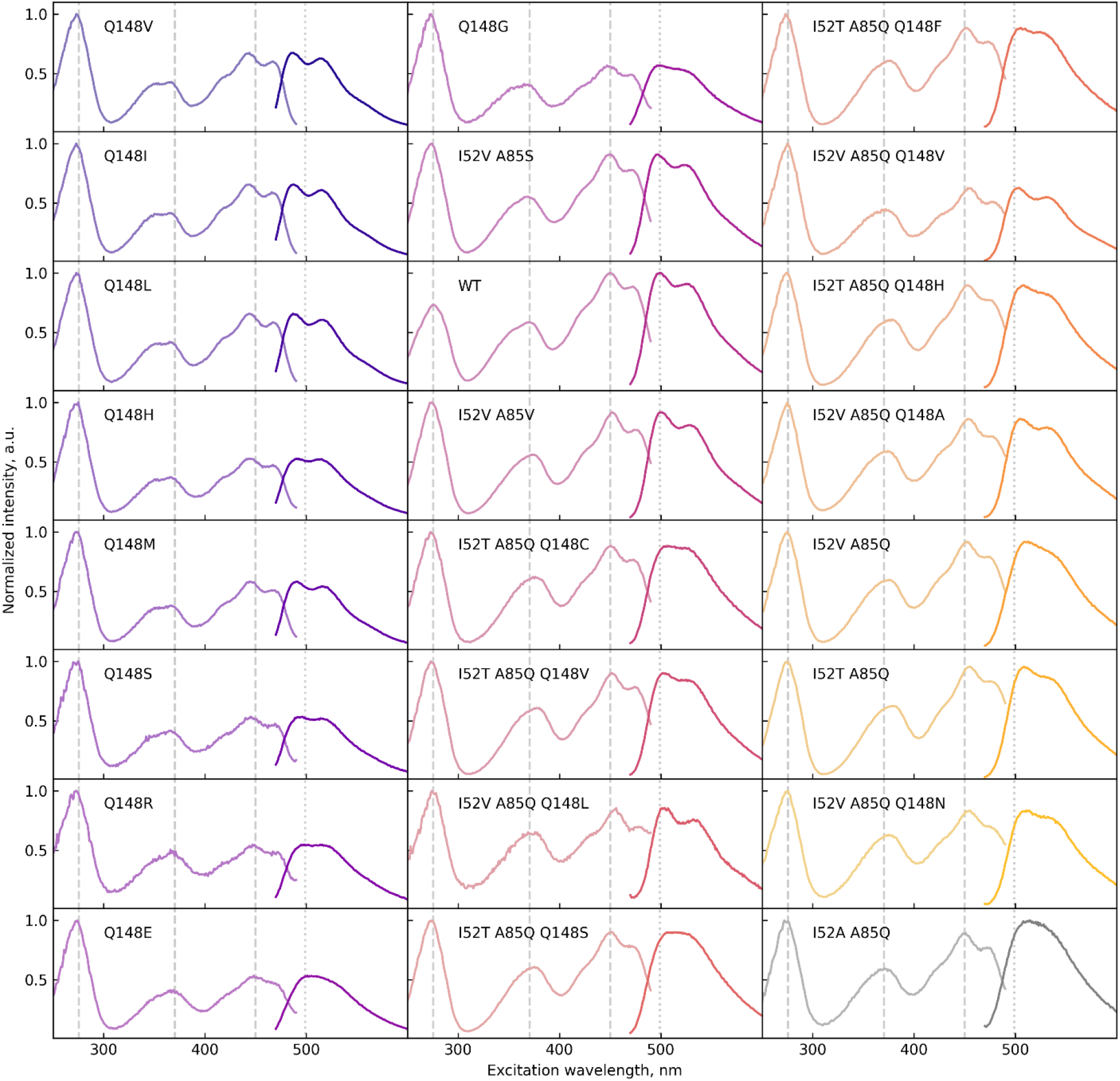
Fluorescence excitation (light colors) and emission (solid colors) spectra of the developed CagFbFP variants. The vertical gray dashed lines show the positions of the main excitation and emission maxima of the parent protein CagFbFP (WT). The I52A A85Q variant (dark gray) is not included in the final palette.

### Fluorescent microscopy using CagFbFP variants

Previously, we reported proof-of-concept two-color fluorescence microscopy imaging with CagFbFP Q148K and CagFbFP I52T Q148K (41). However, these two variants displayed low stability and low expression levels, making them unreliable for applications. Consequently, we decided to test whether the better variants, identified in this work, may be used for two-or three-color microscopy. We selected three mutants from the palette (Q148L, I52T A85Q and the parent CagFbFP) based on their fluorescence emission maxima, expression levels and thermal stability, and expressed them independently in three *E. coli* cultures. The expression conditions were chosen so that the final fluorescence normalized to optical density at 600 nm was within 20% for the three cultures, making it impossible to distinguish the cells based on fluorescence intensity.

We recorded the fluorescence emission of monocultures as well as of the mixture of cells using twelve 9 nm channels in the range from 459 to 557 nm. Spectra from single variant cultures were used as a reference, and a linear unmixing procedure was used to determine the contribution of each of the reference spectra to the fluorescence emission of each pixel in the image of the mixture of cultures (Figure 5). The cells harboring different variants could easily be distinguished from each other, which proves that three-color microscopy using FbFPs is possible.

**Figure 5.**
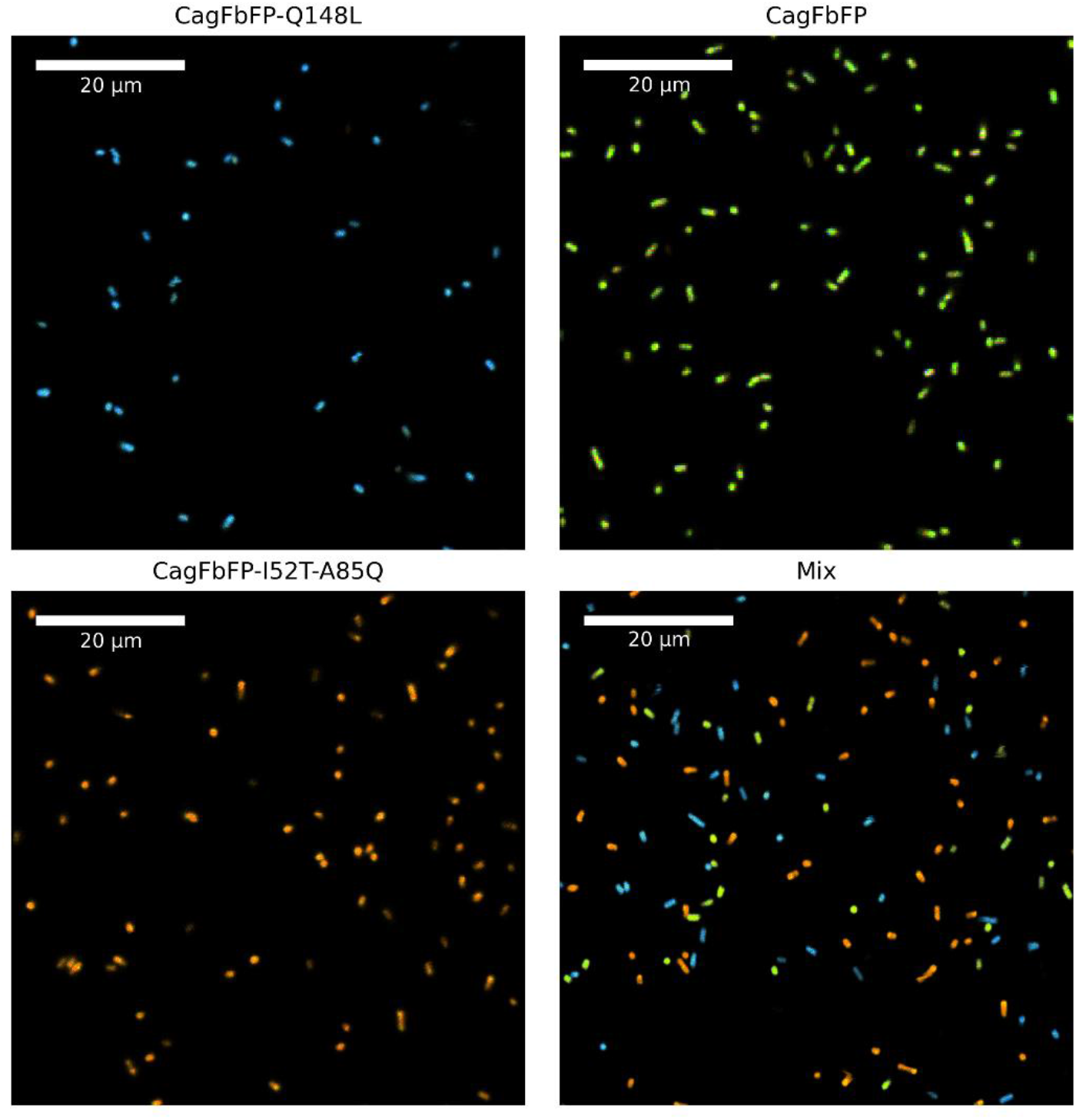
Fluorescence microscopy imaging of *E. coli* cells expressing three different CagFbFP variants. All images were obtained using the same imaging and processing protocols. Single variant cultures were used to obtain reference spectra. Linear unmixing procedure was used to determine the contribution of each of the reference spectra to the fluorescence emission of each pixel, which was then colored accordingly. The red shifted variant is shown in orange, blue shifted is in blue, and parent CagFbFP is in green. The three CagFbFP variants can be unambiguously separated from each other, allowing three-color imaging. The proteins expressed reproducibly, and results of a single experiment are shown in the figure.

To demonstrate that the CagFbFP variants can also be distinguished when present in the same cell, as a proof of concept, we imaged HEK293T cells expressing CagFbFP in mitochondria and CagFbFP-Q148V in the cytoplasm. After linear unmixing, mitochondria can clearly be distinguished from the rest of the cell (Figure 6).

**Figure 6.**
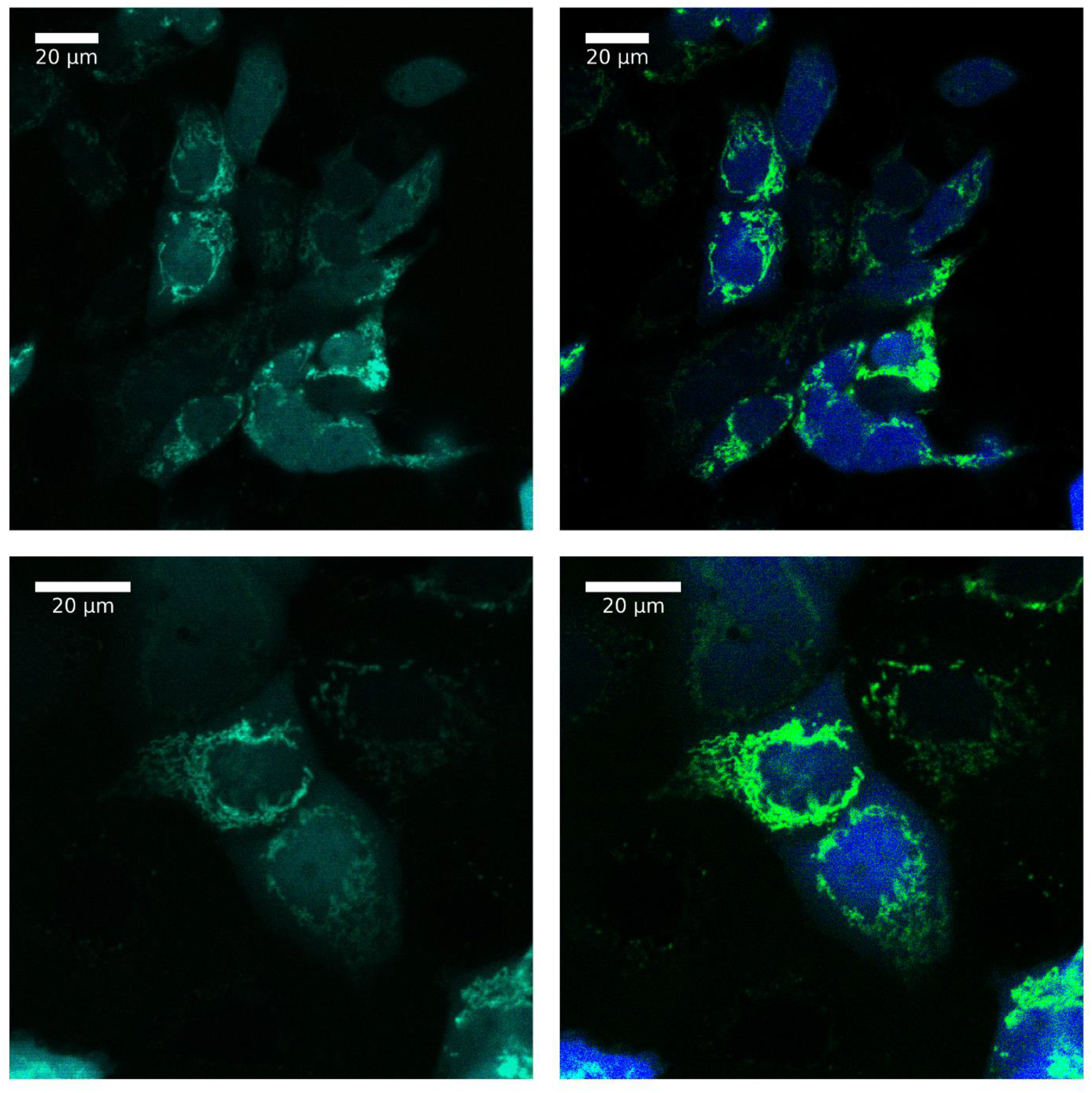
Fluorescence microscopy imaging of HEK293T cells expressing CagFbFP in mitochondria and CagFbFP-Q148V in the cytoplasm. (left) Original images. (right) Spectrally unmixed images. CagFbFP is shown in light green, CagFbFP-Q148V is shown in blue. The two CagFbFP variants can be unambiguously separated from each other, allowing two-color imaging in eukaryotic cells. Cotransfection of HEK293T cells was optimized in a series of 3 experiments, and results of a single experiment are shown.

We also tested whether CagFbFP variants could be separated using fluorescence lifetime imaging microscopy (FLIM) *in vivo.* We expressed the variants Q148H and I52T A85Q, which differ both in emission maxima and in fluorescence lifetimes, in *E. coli* and obtained the microscopy images. The cultures could be distinguished based either on spectra or on fluorescence lifetime (Figure 7), thus confirming the possibility of two-color FLIM using FbFPs. However, we note that the fluorescence lifetimes measured *in vivo* (~2.95 ns and 3.75 ns) differed from those measured *in vitro* (3.24 ns and 4.65 ns), and displayed relatively high cell-to-cell variation; spectrum-based separation is less ambiguous than fluorescence lifetime-based separation.

**Figure 7.**
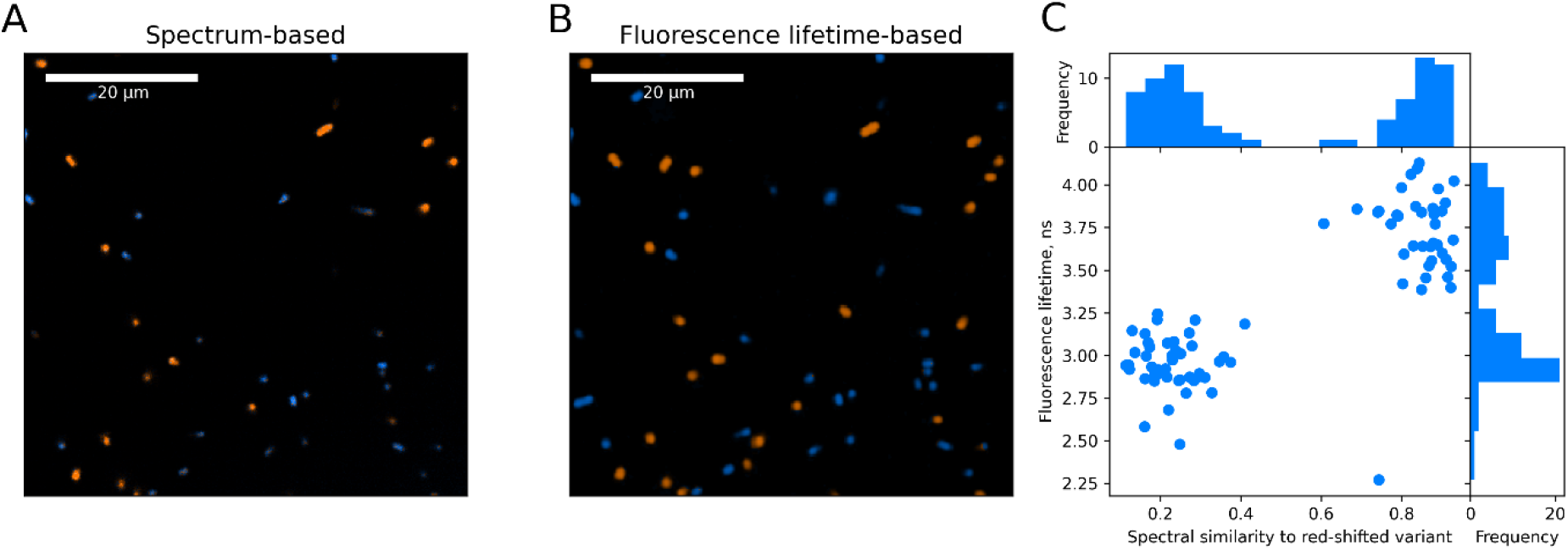
Fluorescence lifetime imaging of *E. coli* cells expressing CagFbFP variants Q148H and I52T A85Q. Same region was imaged using spectral unmixing (A) or fluorescence lifetime measurements (B; pixels displaying fluorescence lifetime below 3.25 ns are shown in blue, and those with lifetime above 3.25 ns are shown in orange). Additional cells entered the field of view in the center image due to prolonged imaging. (C) Quantitative comparison of imaging based on spectral unmixing and fluorescence lifetime measurements. Data for each cell is averaged and shown as a dot. Both procedures give correlated results, with linear unmixing providing better resolution between the two cell types. The proteins expressed reproducibly, and results of a single experiment are shown in the figure.

To further characterize the four CagFbFP variants employed for multicolor imaging, we determined their quantum yields and brightness of fluorescence (Table 2). We note that all of the variants have impaired fluorescence properties compared to the original protein, yet this doesn’t preclude their applications (Figures 5–7).

**Table 2.** Optical properties of selected CagFbFP variants.

| Mutations | Extinction coefficient,<br>$\text{M}^{-1} \text{cm}^{-1}$ | Quantum yield | Brightness, $\text{M}^{-1} \text{cm}^{-1}$ |
| --- | --- | --- | --- |
| Original protein | 15300 | 0.34 | 5200 |
| Q148V | 15400 | 0.33 | 5100 |
| Q148L | 15200 | 0.30 | 4600 |
| Q148H | 14700 | 0.25 | 3700 |
| I52T A85Q | 14000 | 0.22 | 3100 |

## Discussion

Previously, several attempts have been made to generate flavin-based fluorescent proteins with shifted spectra, with moderate success (33–43). In this work, we mutated a previously untargeted amino acid position, that of photoactive cysteine (corresponding to A85 in CagFbFP), in addition to mutations of other residues near the flavin (I52 and Q148, Figure 3A), and identified a palette of 22 variants that uniformly cover the range of emission spectra with the maxima from 486 to 512 nm (Figure 3C). Mutation A85Q produced a notable red shift, but destabilized the protein and thus required compensatory mutations I52T or I52V. The effects of mutation A85Q appear to be roughly additive with the effects of substitutions of Q148. In particular, the mutation Q148V resulted in the largest hypsochromic shifts of the emission maxima for both CagFbFP and CagFbFP I52V A85Q. Interestingly, the variant I52V A85Q Q148V has a spectrum shape almost identical to unmutated CagFbFP, whereas that of I52V A85Q is significantly deformed. While previously only two-color imaging of bacteria using flavin-binding fluorescent proteins was reported (41), here we demonstrate both two-color imaging of eukaryotic HEK293T cells (Figure 6) and three-color imaging of bacteria (Figure 5) employing the new variants.

When expressed in *E. coli*, CagFbFP displays two melting transitions, corresponding to the protein molecules bound to FMN, and those bound to RF/FAD (20). Most mutants identified in this study also show two transitions. Variant Q148H, similarly to some other engineered variants (46), displays three transitions. Some of the mutants are less stable and show only one transition that presumably corresponds to FMN-bound species, thus highlighting the utility of mutating an initially thermostable protein. Except for Q148H, all of the proteins refolded easily after thermal denaturation. We note that all of the red-shifted variants are significantly less stable compared to the original CagFbFP and have lower expression levels, whereas some of the blue-shifted variants (Q148V, Q148I and Q148L) are stabilized and better expressed.

Lifetime of fluorescence for the identified variants ranged from 3.24 ns for the Q148H variant to 4.68 ns for the I52V A85V variant, with no clear correlation with spectral properties (Table 1). The I52T A85Q variant has close to the highest lifetime of 4.65 ns, yet its emission spectrum is significantly different from that of Q148H, which has the lowest fluorescence lifetime. Thus, the pair Q148H and I52T A85Q could potentially be used for two-color imaging based on both spectral separation and fluorescence lifetime. Indeed, we show that two-color fluorescence lifetime-based imaging using CagFbFP variants is possible, although spectral unmixing results in better separation (Figure 7). We note that fluorescence lifetime values as low as 3.17 ns and as high as 5.70 ns were previously reported for other FbFPs; probably, they can provide better separation (34).

## Conclusions

In this work, we showed that the range of color-tuned flavoproteins can be expanded to 25 nm by mutating the previously untested position, that of the photoactive cysteine of FbFP. The position of the emission maximum can also be finely tuned by mutating the conserved glutamine. The developed palette of fluorescent proteins, differing by one, two or three amino acids, can be used for further understanding of the mechanisms of flavin color tuning (47). The obtained variants can be used for three-color imaging based on spectral separation and two-color fluorescence lifetime imaging, paving the way for further applications of FbFPs in fluorescence microscopy.

## Methods

Expression, purification and determination of basic properties follow the procedures outlined in Ref. (48). Determination of advanced properties and fluorescence microscopy imaging follow the procedures outlined in Ref. (49).

### Random mutagenesis and cloning

Site saturation mutagenesis was performed using polymerase chain reaction starting from the original pET11-CagFbFP plasmid (45). For positions 85 and 148, the primers contained a randomized NNK/MNN codon at the target site, surrounded by two complementary ends about 20 nucleotides long, and annealing temperature was set at about 60 °C. The forward and reverse primers were complementary to each other along their entire length, with the exception of the target site. The product of the reaction was a mixture of linear DNA fragments carrying random mutations at the target site. It was transformed into the *E. coli* strain C41 (DE3), where it became circular due to recombination. For position 52, the forward primer was designed without overlap with the reverse primer. The product of the reaction was linear DNA, which was additionally ligated before transformation. The sequences of the primers used for mutagenesis were as follows: I52_forward, CAGATCAGCCGATTGTTTTTG;I52_reverse, CACCGGCATCGGTAAC**MNN**CATACCGCTGGCCATATG;A85_forward, GAATGAAGTTCTGGGTCGTAAT**NNK**CGTTTTCTGCAAGGTCCG;A85_reverse, CGGACCTTGCAGAAAACG**MNN**ATTACGACCCAGAACTTCATTC;Q148_forward, GTTGTTGCATTTGTTGGTGTT**NNK**ACAGATGTTACCGCACATC;Q148_reverse, GATGTGCGGTAACATCTGT**MNN**AACACCAACAAATGCAACAAC.

For fluorescence microscopy of mammalian cells, the CagFbFP gene was fused with N-terminal mitochondrial targeting sequence (MTS, amino acid sequence: MSVLTPLLLRGLTGSARRLPVPRAKIHSLG) as described previously (46). CagFbFP-Q148V gene was cloned into pcDNA3.1(+) via HindIII and BamHI restriction sites.

### Clone selection

The transformed cells were plated on a Petri dish made of LB-agar supplemented with 1 mM IPTG, 150 μg/ml Ampicillin and incubated for 16 hours at 37 °C. After that, 80-90 grown fluorescent colonies were transferred to a 96-well plate (Greiner, Germany), and resuspended in 100 μl of buffer A each (500 mM NaCl, 50 mM Tris-HCl pH 8.0). Emission spectra of each colony were recorded using Synergy H4 Hybrid Microplate Reader (BioTek, USA). The resulting set of spectra was analyzed for brightness and spectral shift. Based on spectral data from all the mutagenesis trials performed, the final set of 22 mutants was selected for the consideration of uniform coverage of the widest possible spectral shift range with a good expression level and thermal stability.

### Expression and purification

Selected colonies containing plasmids with genes of desired protein were sequenced and cultured in shaking flasks in 200-250 ml LB medium containing 150 mg/l ampicillin. Protein expression was induced by addition of 1mM IPTG and followed by incubation for 24 hours at 16 °C. Harvested cells resuspended in lysis buffer A were disrupted by incubation in a water bath at 95 °C for 10 minutes with an additional 10 minutes incubation at 0 °C for protein refolding. Cell debris was removed by centrifugation at 15000 g for 30 minutes at 10 °C.

The obtained supernatant was incubated with Ni-nitrilotriacetic acid (Ni-NTA) resin (Qiagen, Germany) in a rotating shaker for 1 h. The resin (separated from supernatant by centrifugation at 200 g for 3 minutes) was resuspended in 20 column volumes of buffer A and again separated by centrifugation for washing. For a better result, this procedure was performed at least two times.

Cleaned Ni-NTA resin with bound protein was incubated in 5 column volumes of buffer A with addition of 200 mM imidazole at room temperature for 5 minutes and separated by centrifugation at 200 g for 3 minutes. The supernatant containing most of the resulting protein was collected in a separate tube. To collect the residue protein bound to the resin this procedure was repeated with 400 mM of imidazole. The two fractions were transferred to the final buffer (100 mM NaCl, 20 mM Tris-HCl pH 8.0) by dialysis.

### Spectroscopy

*In vivo* and *in vitro* spectra of all proteins were recorded using Synergy H4 Hybrid Microplate Reader (BioTek) with resolution of 9 nm. Emission spectra were measured between 470 and 700 nm with excitation at 450 ± 5 nm. Raw spectra were fitted using a sum of four Gaussians defined as a function of the wavenumber, and a constant background. Positions of the maxima λem and of the Gaussians corresponding to the first and the second peaks of the spectra were determined from the fit.

For calculation of the hue parameter H, raw spectra were used. Each spectrum was initially converted to the XYZ color representation using CIE 1931 color space, and after that to HSV color representation using the colormath Python library by Greg Taylor (https://github.com/gtaylor/python-colormath). H is measured in degrees, ranging from 0 to 360.

### Thermal stability

Melting curves of proteins were measured using the Rotor-Gene Q real-time PCR cycler (Qiagen, Germany). 25 μl samples with a protein concentration of approximately 0.5 mg/ml in a buffer containing 100 mM NaCl, 20 mM Tris-HCl pH 8.0, were used. Fluorescence was excited at 470 ± 10 nm, and emission was recorded at 510 ± 5 nm. Samples were heated up from 30 °C to 95 °C at a heating rate of 1 °C per minute and held at 95 °C for 5 minutes. After that they were cooled down to 30 °C with a cooling rate of 1 °C per minute and held for 5 minutes at 30 °C to maximize refolding. During the whole procedure, fluorescence intensity was measured as a function of temperature with 0.5 °C step. Data was denoised using the Savitzky–Golay filter. First derivatives were taken using symmetric difference quotient. Since each derivative of the melting curves could consist of two or more peaks, we applied multi-Gaussian fitting with a constant baseline to obtain more accurate positions of peaks.

### Determination of quantum yields

Given that QY of CagFbFP depends on the chromophore composition, which varies with expression conditions (20), all samples were refolded with FMN prior to measurements. The protein in the apo form prepared as described in ref. (20) was mixed with 4 mM FMN (Sigma-Aldrich, USA) to reach 3:5 FMN:protein molar ratio. After incubation for 10 min, the samples were transferred into the buffer containing 50 mM Tris pH 8.0, 300 mM NaCl by dialysis.

Samples were diluted to optical density ≤ 0.1 in order to minimize inner filter effects. For emission measurements, samples were excited by a M455F3 LED at 455 nm (Thorlabs, USA), and the emission spectra were recorded between 457 and 648 nm using AvaSpec-2048L spectrometer (Avantes, Netherlands). For absorbance measurements, samples were illuminated by AvaLight-DHc full-range light source (Avantes, Netherlands), and spectra were recorded by the same AvaSpec-2048L spectrometer. The temperature of the samples was maintained at 20 °C by qpod 2e temperature-controlled sample compartment (Quantum Northwest, USA). QY was determined relative to the original CagFbFP: 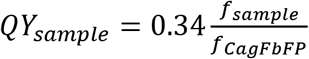, with 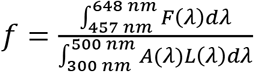, where F and A are emission and absorption spectra of the sample and L is the emission spectrum of LED.

### Determination of the extinction coefficients

The samples were reconstituted with FMN as described in the previous section. Spectroscopic measurements were carried out using the same setup. The absorption spectrum of the solution of FMN was obtained at 20°C and at 95°C. Using the known value of the extinction coefficient of flavins at room temperature (12020 M^-1^ cm^-1^ at 445 nm), the extinction coefficient of the chromophore solution at 95°C was determined to be equal 11550 M^-1^ cm^-1^. The absorption spectrum of the sample was measured at 20 °C, then the sample was heated to 95 °C. During the heating, protein aggregation occurred, with a rapid increase in OD. At approximately 90 °C, the aggregates dissociated and the absorption spectrum of the sample corresponded to that of the free chromophore. Using this spectrum, the molar concentration of the chromophore in the sample was measured. The protein extinction coefficient was determined as the ratio of the maximum absorption in the 400-500 nm range to the molar concentration of the chromophore in the assumption that all of the chromophore in the solution was bound to the protein at 20 °C.

### HEK293T cell culture and transfection

HEK293T cells were grown in Dulbecco’s modification of Eagle’s medium (DMEM) supplemented with 10% heat-inactivated fetal bovine serum, 1% Penicillin/Streptomycin, 2 mM GlutaMax and 10 mM HEPES (all from Thermo Fisher Scientific, USA) at 37 °C and 5% CO_2_. The cells were plated in a 35 mm dish with glass bottom (ibidi, Germany) in full medium 48 h before transfection. The cells were transfected using Lipofectamine LTX Reagent with PLUS Reagent (Thermo Fisher Scientific, USA) according to the manufacturer’s instructions. For cotransfection, 2 μg total plasmid DNA (725 ng CagFbFP-Q148V plasmid + 1275 mg MTS-CagFbFP plasmid) and 2 μl PLUS Reagent were diluted in 100 μl Opti-MEM (Thermo Fisher Scientific, USA), and 4 μl Lipofectamine LTX were diluted in 100 μl Opti-MEM. Diluted DNA was added to diluted Lipofectamine LTX Reagent, the mixture was incubated for 10 min and added to the cells in full medium. The cells were examined 20-24 h after transfection.

### Fluorescence microscopy of E. coli and HEK293T cells

Fluorescence confocal laser scanning microscopy (CLSM) was performed on an inverted fluorescence microscope LSM780 (Zeiss, Germany), using oil immersion 100× objective (NA=1.46, Zeiss) for *E. coli* cells and oil immersion 63× objective (NA=1.4, Zeiss) for mammalian cells. Solutions of *E. coli* cells were diluted to the OD_600_ of ~0.1 and placed into the 8 chambered glass bottom microscopy slide (ibidi, Germany). Live HEK293T cells were maintained at 37 °C and 5% CO_2_ in INUBG2H-ELY incubator (TOKAI HIT, Japan) during the measurements.

Laser excitation of the fluorescence was done using Ar-Ion laser (Lasos, Germany) at 458 nm wavelength. Fluorescence images were obtained by a 34-channel QUASAR detector (Zeiss, Germany) set to the 460-690 nm wavelength range. Fluorescence spectra for *E. coli* cultures expressing different mutants were used to provide reference spectra for the spectral unmixing of the mixture of cells expressing different mutants. The same approach was used for HEK293T. Spectral unmixing was performed using the *fnnls* Python library by Joshua Vendrow and Jamie Haddock (https://pypi.org/project/fnnls/) (50).

### Fluorescence lifetime measurements and microscopy

Fluorescence lifetime measurements and fluorescence lifetime imaging microscopy (FLIM) were performed on an inverted fluorescence microscope LSM780 (Zeiss, Germany), using oil immersion 63× objective (NA=1.4, Zeiss). Solutions of *E. coli* cells were diluted to the OD_600_ of ~0.1 and placed into the 8 chambered glass bottom microscopy slide (ibidi, Germany). Two photon excitation was achieved using Insight DeepSee laser (λ_exc_ = 920 nm; 80 MHz, 150 fs pulse; Newport Spectra Physics, USA). Fluorescence intensity decays were generated using time-correlated single photon counting (TCSPC) with SPC-150 modules (Becker &; Hickl, Germany). Average lifetimes were calculated using monoexponential fitting with a binning parameter of 3 in SPCImage software (Becker &; Hickl), and averaged to obtain τ_av_. CagFbFP solutions with concentration of ~0.1 mg/ml were measured using the same experimental setup.

## Funding

The study was funded by the Russian Science Foundation, grant number 21-64-00018.

## Notes

### Competing Interest Statement

The authors have declared no competing interest.

